# Characteristics of University Students Supported by Counseling Services: Analysis of psychological tests and pulse rate variability

**DOI:** 10.1101/658740

**Authors:** Yumi Adachi, Hiroaki Yoshikawa, Shigeru Yokoyama, Kazuo Iwasa

## Abstract

**Objective:** The number of students who use counseling services is increasing in universities. We clarified the characteristics of the students who access counseling services using both psychological tests and pulse rate variability (PRV).

**Methods:** We recruited the participants for this study from the students who had counseling sessions at Kanazawa University (Group S). As a control group, we also recruited students who had no experience in counseling sessions (Group H). We obtained health information from the database of annual health check-ups. Participants received the Wechsler Adult Intelligence Scale (WAIS) III, Autism-Spectrum Quotient (AQ), Sukemune-Hiew (S-H) Resilience Test, and State-Trait Anxiety Inventory-JYZ (STAI). We also studied the SF-12v2 Health Survey. We examined the spectral analyses of pulse rate variability (PRV) by accelerating plethysmography. We performed linear analysis of PRV for LF, HF, and LF/HF as indexes of autonomic nervous function. We also performed non-linear analysis of PRV for the largest Lyapunov exponent (LLE). We examined participants’ blood for autoantibodies against glutamate decarboxylase (GAD) 65.

**Results:** A total of 105 students participated in this study. Group S had 37 participants (Male: 26, Female: 11) and Group H had 68 participants (Male: 27, Female 41). There were 12 males and two females who had been diagnosed with autism spectrum disorder (ASD) in Group S. Group S had 1) lower Working Memory Index (WMI) despite relatively high Full-Scale IQ, 2) higher AQ scores, 3) lower resilience scores, 4) higher anxiety scores, 5)lower QOL in social health, and 6) shifting of autonomic nervous balance toward higher sympathetic activity. There were only two participants who had the anti-GAD65 antibody in Group S, and one participant had the anti-GAD65 antibody in Group H.

**Conclusion:** The results of our study revealed valuable information about the students who seek mental support in our university. Our results provide information that can be used in interventions for the university students who need counseling services.

## Introduction

During adolescence, mental health is an important issue [1], and one of the leading causes of premature death in young people is suicide [2]. There have been many attempts to prevent suicide in adolescents [3]. The numbers of students seeking counseling services are increasing in universities [4]. The complaints of students who come to counseling services are varied; however, in a survey in the United States, the major factors preventing students from academic success are: anxiety disorders (89%), crises requiring immediate response (69%), psychiatric medication issues (60%), clinical depression (58%), learning disabilities (47%), sexual assault on campus (43%), self-injury issues (e.g. cutting to relieve anxiety) (35%), and problems related to earlier sexual abuse (34%) [5]. Hikikomori (acute social withdrawal) is a severe mental disability that affects students in Japan [6]. It is difficult to treat sufferers once they show the symptoms of Hikikomori. Recently, patients with Hikikomori have been diagnosed by psychiatrists in a variety of cultures and countries [7]. These conditions are often associated with developmental disorders including autism spectrum disorder (ASD) [8]. Recently, a considerable number of students with ASD traits, who were not diagnosed with ASD, enter universities. Their difficulties with socialization have been found to affect their academic success and overall well-being [9]. Occasionally, university life reveals students’ mild developmental disorders such as when students live alone apart from their parents. ASD has many symptoms in addition to difficulties in communication. For example, hypersensitivity to sensory stimuli especially auditory stimuli, bowel problems, and hyperhidrosis are common. These symptoms may be caused by impairments of autonomic nervous functions [10].

In the counseling rooms of universities, licensed counselors have sessions with students. Counselors assess clients’ conditions from a psychological perspective. The most common issues are the students’ complaints with or difficulties in university life, which are usually reported by subjective dialogs only with psychological tests not necessarily being performed. Patients often complain about physical symptoms including insomnia, hyperhidrosis and irritable bowels, which may be related to autonomic nervous system dysfunction. A large number of students are referred to psychiatric services following counseling and receive medical diagnoses. In the case of Hikikomori, diagnosis is difficult. In this study, we attempted to evaluate the mental and physical conditions of students who had counseling sessions by comparing them with students who did not use counseling services. To assess their individual states, we performed a series of psychological tests. We also tested autonomic nervous functions by spectral analyses of pulse rate variability (PRV) measured by accelerating plethysmography. Non-linear analyses of the pulse wave (largest Lyapunov exponent, LLE) recorded from the subjects’ fingertips provide results related to the central nervous system [11]. As a biological marker for ASD and ADHD, we tested for the presence of auto-antibodies against glutamate decarboxylase (GAD65), which Rout *et al.* detected in 15% of ASD and 27% of ADHD children [12]. From these examinations, we attempted to identify the specific characteristics of students who sought counseling sessions in the university.

## Materials and Methods

### Study design

This study is part of a randomized, cross-over, placebo-controlled trial (http://www.umin.ac.jp/ctr/index.htm, number UMIN000019101) in a single center (Kanazawa University, Kanazawa, Japan). Participants were recruited from students of Kanazawa University. The study was conducted with Good Clinical Practice. Enrolment started in October 2015, and the last participant finished their observational period in October 2016. This report is a summary of the pre-evaluation of participants before entering the placebo-controlled trial.

### Standard protocol approvals, registrations, and participant consents

The study was approved by the Ethics Committee of Medicine, Kanazawa University (No. 29-3), and written informed consent was obtained from all subjects enrolled.

### Participants

We recruited participants from students of Kanazawa University (https://www.kanazawa-u.ac.jp/e/). Group S consisted of students who had counseling sessions in our counseling rooms. Group H consisted of students, recruited from classes, who had no experience in counseling sessions. As exclusion criteria, students who had severe mental disorders or a suicide attempt were excluded from this study.

### Assessment

We interviewed candidates and screened their eligibility for this study. We obtained full informed consent using printed materials. We accessed health information on the participants from the database of annual health check-ups in the Health Service Center, Kanazawa University. Participants received the Wechsler Adult Intelligence Scale (WAIS) - Third Edition (III), Autism-Spectrum Quotient (AQ), Sukemune-Hiew (S-H) Resilience Test and State-Trait Anxiety Inventory-JYZ (STAI). We also studied the SF-12v2 Health Survey (iHope International Co. Ltd., Kyoto, Japan). To evaluate the function of the autonomic nervous system, we investigated the spectral analyses of PRV measured by accelerating plethysmography. We also performed non-linear analysis of the data from the pulse wave study and obtained the values of the largest Lyapunov exponent (LLE). To study autoantibodies against GAD65, we drew 5 mL of blood from participants. The sera was separated and then stored at −80°C until antibody testing. The order of the assessments was as follows: 1) SF-12v2, 2) STAI, 3) S-H Resilience Test, 4) AQ, 5) PRV, 6)blood drawing, and 7) WAIS-III. The WAIS-III was performed on a different day from the other tests because of time constraints.

### WAIS-III

The version of WAIS-III used in this experiment was the Japanese version purchased from Nihon Bunka Kagakusha (Tokyo, Japan). The original version is published by Pearson (USA). The test was performed by licensed psychocounselors.

### Autism-Spectrum Quotient (AQ)

The definition of AQ used in this experiment was based on the Japanese version of the original by Baron-Cohen *et al.* [13], which was revised in 2016. AQ has five subcategories: 1) Social skill (for example: “I prefer to do things with others rather than on my own.”—reversal item); 2) Attention switching (for example: “I prefer to do things the same way over and over again”); 3) Attention to detail (for example: “I often notice small sounds when others do not.”); 4) Communication (for example: “Other people frequently tell me that what I’ve said is impolite, even though I think it is polite.”); 5) Imagination (for example: “When I’m reading a story, I can easily imagine what the characters might look like”—reversal item). The maximum score for each subcategory is 10. The total score is out of 50 and ASD is suspected when the score is equal to or more than 33.

### Sukemune-Hiew (S-H) Resilience Test

Sukemune-Hiew (S-H) Resilience Test was developed and validated by Sato and Sukemune [14] to evaluate the power of resilience in adults. We purchased printed test sheets from Takei Scientific Instruments Co., Ltd. (Niigata, Japan).

### State-Trait Anxiety Inventory-JYZ (STAI)

The State-Trait Anxiety Inventory (STAI) was initially developed by Spielberger *et al.* [15], and the Japanese version (STAI-JYZ) was made by Hidano *et al.* (Jitsumu Kyoiku Shuppan, 2000). We purchased printed test sheets from Jitsumu Kyoiku Shuppan (Tokyo, Japan).

### SF-12v2 Health Survey

To evaluate health-related quality of life (QOL), we used the SF-12v2 Health Survey (iHope International Co. Ltd., Kyoto, Japan). We evaluated lower measures of SF-12v2, which were composed of Physical Functioning (PF), Role Physical (RP), Bodily Pain (BP), General Health (GH), Vitality (VT), Social Functioning (SF), Role Emotional (RE), and Mental Health (MH). Then we transformed the scores from these summaries into Physical component summary (PCS), Mental component summary (MCS) and Role/Social component summary (RCS) scores [16].

### Spectral analyses of PRV measured by acceleration plethysmography and LLE by non-linear analysis of PRV

We performed spectral analyses of PRV measured by acceleration plethysmography. We collected data with an Android™ tablet installed with Alys™ (Chaos Technology Research Laboratory, Otsu, Japan) at a stable temperature and under quiet conditions. Participants sat on chairs and were asked to relax for five minutes before recording. We used fingertip sensors to record heart rates. If the recorded waves were too small to analyze correctly, we used sensors attached to the subjects’ earlobes. We recorded the data with the subject in a sitting position for three minutes. The collected data was calculated using Lyspect™ (Chaos Technology Research Laboratory, Otsu, Japan) installed in a PC with Windows 10. For spectral analyses, we used fast Fourier transform (FFT). We set a low-frequency (LF) range from 0.04 to 0.15 Hz, and a high-frequency (HF) range from 0.15 to 0.40 Hz. The densities of the power spectrum were calculated from LF and HF, respectively, and the power ratio of LF/HF was obtained, which was estimated for the sympathetic activities. As an index of parasympathetic activities, we used the power spectrum of HF. We also performed non-linear analysis of the pulse wave results and obtained LLE from the same data for the power spectrum.

### Anti-GAD65 antibody

To measure anti-GAD65 antibody levels in serum, we used GADAb ELISA kits (Cosmic Corporation, Tokyo, Japan). Participants’ sera were tested using the instructions provided by the vendor.

### Statistical methods

For statistical analyses, we applied t-test to the data which had a normal distribution and Wilcoxon signed rank test to the data which had a non-normal distribution. We analyzed data using IBM SPSS Statistics ver. 22 (IBM Japan, Tokyo, Japan).

## Results

### Participant disposition

A total of 105 participants were enrolled in this study. Participants’ age and health information were obtained from annual health checkups performed in April (Table 1). In both males and females, age was higher in Group S than Group H. In the male subjects, age and body weight (BW) were higher in Group S than Group H. In the female subjects, age was higher in the subjects in Group S. In Group S, 12 male participants and two female participants had been diagnosed with ASD.

**Table 1.** Clinical information of participants.

| Male | S (N=26) | H (N=27) |  |
| --- | --- | --- | --- |
|  | median (interquartile range) | median (interquartile range) |  |
| Age | 22 (21-24) | 20 (19-21) | p=0.0003 |
| Height (cm) | 172 (168-178) | 170 (168-174) | p=0.1555 |
| BW (kg) | 62 (61-73) | 57 (56-62) | p=0.0069 |
| BMI | 21.5 (19.3-23.9) | 20.3 (19.5-21.3) | p=0.0781 |
| Clinical Diagnosis | ASD; 12 (46.1%) | none |  |
| Female | S (N=11) | H (N=41) |  |
|  | median (interquartile range) | median (interquartile range) |  |
| Age | 23 (20-27) | 20 (19-20) | p=0.0021 |
| Height (cm) | 161 (157-166) | 161 (157-163) | p=0.9373 |
| BW (kg) | 53 (48-57) | 52 (47-54) | p=0.4723 |
| BMI | 20.4 (18.8-23.6) | 19.9 (18.4-21.6) | p=0.4529 |
| Clinical Diagnosis | ASD: 2 (18.2%) | none |  |

Interquartile range = difference between 25<sup>th</sup> and 75<sup>th</sup> percentiles

### WAIS-III

There was no difference in Full-Scale IQ (FSIQ), Verbal IQ (VIQ) and Performance IQ (PIQ) between Group S and Group H (Table 2). Working Memory Index (WMI) and Processing Speed Index (PSI) were lower in Group S (Table 2).

**Table 2.** WAIS-III IQ and Index.

|  | Group S | Group H |  |
| --- | --- | --- | --- |
| | mean ± SD | mean ± SD | |
| Full-Scale IQ (FSIQ) | 115.7 ± 11.1 | 117.6 ± 8.3 | p=0.309 |
| Verbal IQ (VIQ) | 120.1 ± 10.0 | 121.3 ± 8.7 | p=0.524 |
| Performance IQ (PIQ) | 107.0 ± 15.8 | 108.9 ± 9.9 | p=0.506 |
| Verbal Comprehension Index (VCI) | 121.2 ± 10.8 | 117.5 ± 9.3 | p=0.070 |
| Perceptual Organization Index (POI) | 106.8 ± 15.6 | 108.1 ± 11.7 | p=0.630 |
| Working Memory Index (WMI) | 106.2 ± 12.5 | 111.8 ± 11.7 | p=0.026 |
| Processing Speed Index (PSI) | 103.2 ± 17.7 | 109.1 ± 12.0 | p=0.045 |

In the Verbal tasks, Similarities was higher in Group S. Letter-Number Sequencing in the Verbal tasks and Digit Symbol Search in the Performance tasks were lower in Group S (Table 3).

**Table 3.**
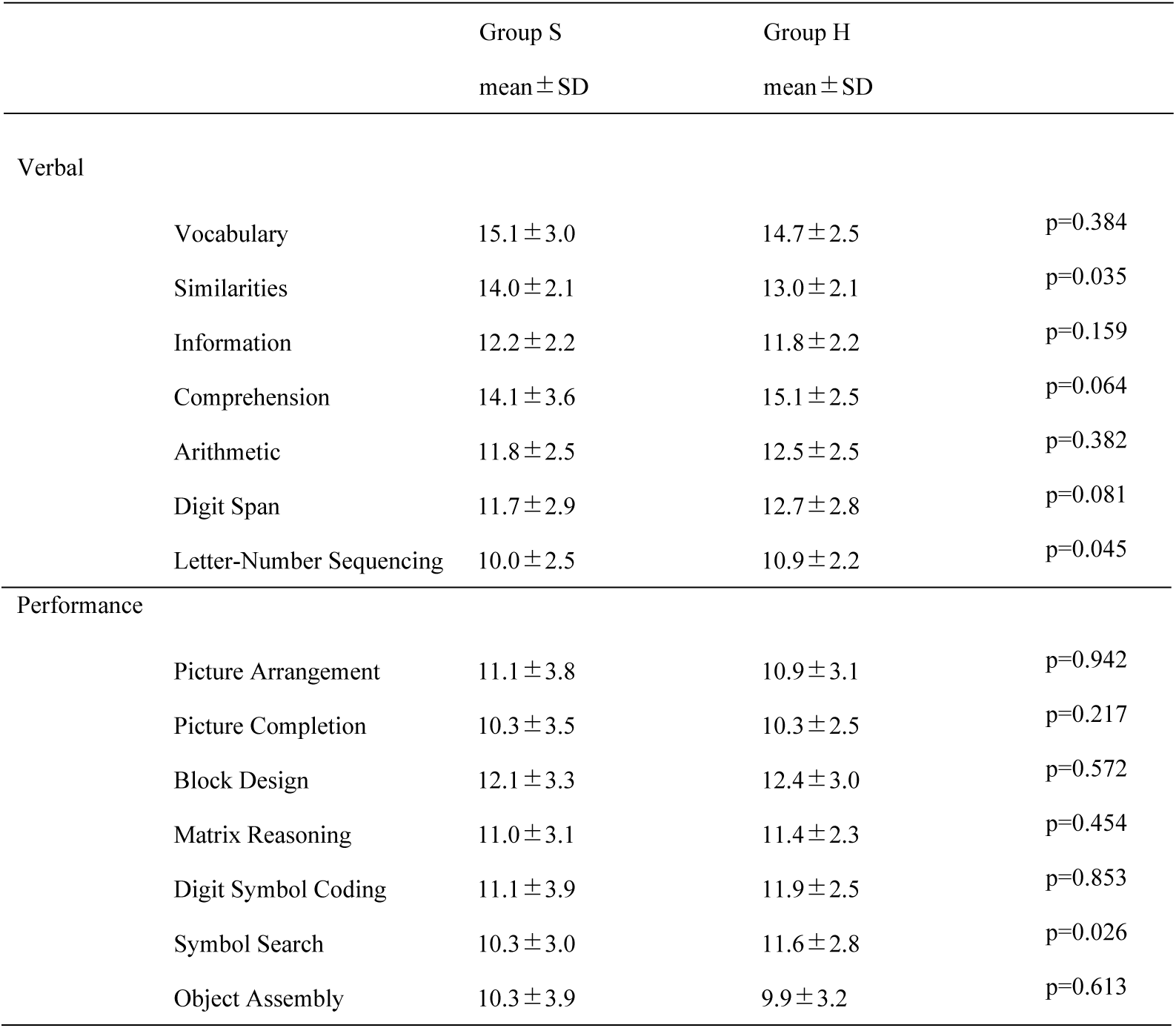
WAIS-III Tasks.

|  | Group S | Group H |
| --- | --- | --- |
| | mean ± SD | mean ± SD |

Verbal
|  |  |  |  |
| --- | --- | --- | --- |
| Vocabulary | 15.1 ± 3.0 | 14.7 ± 2.5 | p=0.384 |
| Similarities | 14.0 ± 2.1 | 13.0 ± 2.1 | p=0.035 |
| Information | 12.2 ± 2.2 | 11.8 ± 2.2 | p=0.159 |
| Comprehension | 14.1 ± 3.6 | 15.1 ± 2.5 | p=0.064 |
| Arithmetic | 11.8 ± 2.5 | 12.5 ± 2.5 | p=0.382 |
| Digit Span | 11.7 ± 2.9 | 12.7 ± 2.8 | p=0.081 |
| Letter-Number Sequencing | 10.0 ± 2.5 | 10.9 ± 2.2 | p=0.045 |

Performance
|  |  |  |  |
| --- | --- | --- | --- |
| Picture Arrangement | 11.1 ± 3.8 | 10.9 ± 3.1 | p=0.942 |
| Picture Completion | 10.3 ± 3.5 | 10.3 ± 2.5 | p=0.217 |
| Block Design | 12.1 ± 3.3 | 12.4 ± 3.0 | p=0.572 |
| Matrix Reasoning | 11.0 ± 3.1 | 11.4 ± 2.3 | p=0.454 |
| Digit Symbol Coding | 11.1 ± 3.9 | 11.9 ± 2.5 | p=0.853 |
| Symbol Search | 10.3 ± 3.0 | 11.6 ± 2.8 | p=0.026 |
| Object Assembly | 10.3 ± 3.9 | 9.9 ± 3.2 | p=0.613 |

### AQ score

Total AQ score was higher in Group S (Table 4). This score indicates that participants in Group S had higher traits for ASD. The number of participants who were suspected of having ASD based on their AQ score (≥33) was nine (24.3%) in Group S and three (4.4%) in Group H. Group S had difficulties in Social skill, Attention switching, Communication, and Imagination. There was no difference in Attention to detail between Group S and Group H.

**Table 4.**
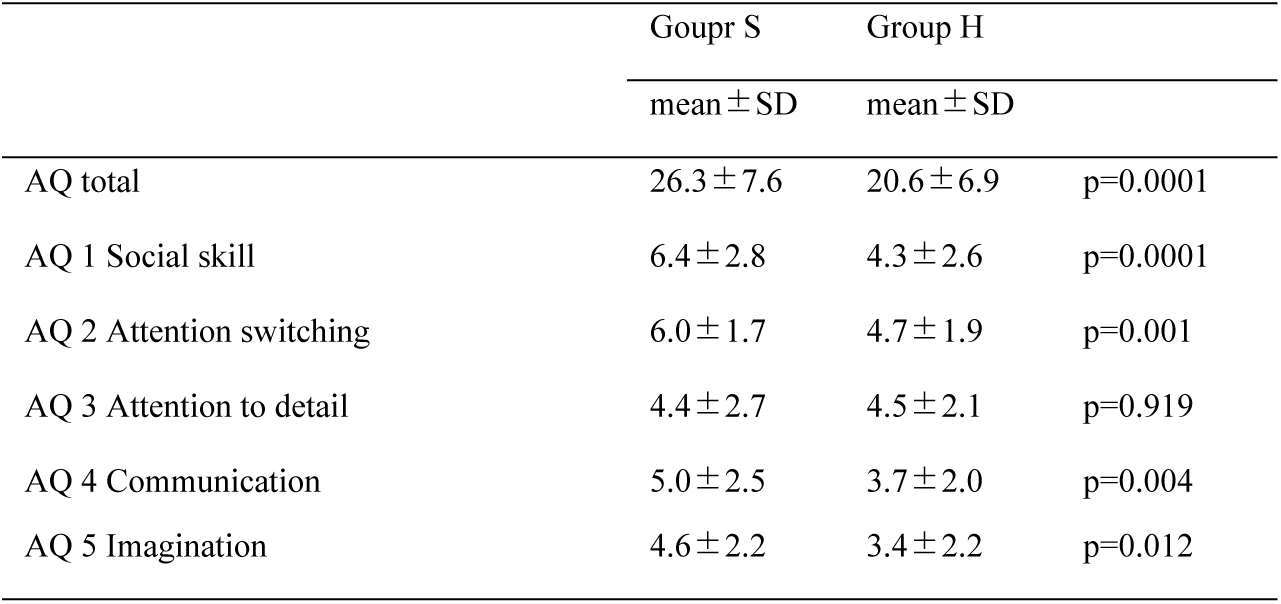
AQ scores of participants.

### The results of the S-H Resilience Test and STAI

The results of the S-H Resilience Test and STAI are summarized in Table 5. S-H Resilience Score was lower in Group S. It means that Group S had less resilience than Group H. As far as STAI, Group S showed higher scores compared with Group H in both State Anxiety and Trait Anxiety.

**Table 5.** S-H Resilience Test and STAI.

|  | Group S (N=37) | Group H (N=68) |  |
| --- | --- | --- | --- |
| | mean ± SD | mean ± SD | |
| S-H Resilience Test | 88.1 ± 10.0 | 102.7 ± 13.1 | p=0.0001 |
| STAI State Anxiety Score | 43.7 ± 11.0 | 39.1 ± 8.8 | p=0.023 |
| STAI Trait Anxiety Score | 54.7 ± 10.6 | 46.2 ± 10.1 | p=0.0001 |

### SF12v2 Health Survey

The results of the SF12v2 Health Survey are summarized in Table 6. Group S had lower scores in Role/Social component summary (RCS).

**Table 6.** SF12v2 Health Survey.

|  | Group S (N=37) | Group H (N=68) |  |
| --- | --- | --- | --- |
|  | median (Interquartile range) | median (Interquartile range) |  |
| Physical component summary (PCS) | 59.0 (52.7-65.0) | 57.5 (54.2-61.4) | p=0.3674 |
| Mental component summary (MCS) | 49.8 (43.9-54.0) | 51.2 (46.6-55.7) | p=0.1826 |
| Role/Social component summary (RCS) | 37.4 (30.1-42.5) | 44.9 (37.7-52.0) | p=0.0009 |

Interquartile range = difference between 25<sup>th</sup> and 75<sup>th</sup> percentiles

### Analysis of pulse waves

Linear and non-linear analyses of the pulse wave results were obtained by accelerating plethysmography. We obtained the powers of LF and HF by spectral analysis of PRV. The ratio of LF/HF is estimated as sympathetic nervous tone, and HF is representative of activities of the parasympathetic nerve. The LF/HF was higher in Group S than Group H. The HF was lower in Group S. This means that the activity of the autonomic nerve of Group S decreases to the hyper sympathetic status (Table 7). Non-linear analysis of PRV brought us the LLE of the attractor, which is constructed for the time series data from pulse waves. There was no difference in LLE and LLE (SD) between Group S and Group H. The mean value of heart rate (HR) and coefficient of variation of R-R intervals (CVRR %) was not different between Group S and Group H (Table 8).

**Table 7.** Pulse wave analysis.

|  | Group S (N=37) | Group H (N=66) |  |
| --- | --- | --- | --- |
|  | median (interquartile range) | median (interquartile range) |  |
| LF | 9.50 (5.12-12.97) | 7.62 (5.14-12.74) | p=0.831 |
| HF | 5.11 (3.12-9.22) | 6.96 (3.76-10.31) | p=0.245 |
| LF/HF | 1.77 (1.08-2.74) | 1.22 (0.65-2.00) | p=0.038 |
| LLE | 4.39 (3.74-6.28) | 5.09 (4.01-6.00) | p=0.379 |
| LLE (SD) | 1.114 (0.77-1.54) | 1.06 (0.85-1.48) | p=0.752 |

Interquartile rage = difference between 25<sup>th</sup> and 75<sup>th</sup> percentiles

**Table 8.** HR and CVRR.

|  | Group S (N=37) | Group H (N=66) |  |
| --- | --- | --- | --- |
| | mean ± SD | mean ± SD | |
| HR (mean) | 74.7 ± 11.3 | 74.1 ± 8.8 | p=0.761 |
| CVRR (%) | 7.16 ± 5.15 | 6.50 ± 2.10 | p=0.459 |

HR = heart rate, CVRR = Coefficient of Variation of R-R intervals

### Anti-GAD65 Antibody

The level of anti-GAD65 antibody was tested using participants’ sera. The cutoff value settled at 5.0 U/mL. There were only three participants who had positive results, two from Group S (9.0 U/mL, 7.7 U/mL) and one from Group H (10.2 U/mL). The two participants with a positive result for anti-GAD65 antibody from Group S had been diagnosed with ADH. There was no overall difference in anti-GAD65 antibody titers between Group S and Group H (Table 9).

**Table 9.** Anti-GAD65 antibody.

| | Group S(N=31)<br>mean ± SD | Group H(N=53)<br>mean ± SD | |
| --- | --- | --- | --- |
| anti-GAD65 antibody (U/mL) | 1.91 ± 2.10 | 1.81 ± 1.87 | p=0.812 |

## Discussion

The purpose of this study was to clarify the commonalities between students who utilize the counseling services of the university. Students of Group S received counseling sessions. The average age of Group S was higher than Group H for both the female and male participants (Table 1). BW was higher in the males of Group S than Group H, but there was no difference in BMI. The higher BW may be attributable to two participants who had a considerable BW. As far as accompanying conditions, 46.1% of males in Group S and 18.2% of females in Group S had been diagnosed with ASD by psychiatrists.

WAIS-III revealed several characteristics of Group S. Working Memory Index was low in Group S (Table 2). Regarding the tasks that were studied, Similarities was high in Group S, and Digit Span and Symbol Search were low in Group S. These results indicate the superiority of verbal function and problems with working memory in Group S. Working memory is a function whereby the person stores useful information in their mind for a short period, which typically decreases with age [17]. It is related to the functional connectivity of large scale brain network [18].

The AQ score was higher in Group S (Table 4). Group S has a higher trait toward ASD by AQ. There were nine participants (24.3%) who had a higher score (≥33) in Group S and three participants in Group H who had a higher score (≥33). From our results, it is evident that students with high AQ scores do not necessarily face difficulties in their academic lives. As far as subcategories, Group S exhibited high scores in every subcategory except for Attention to detail.

Regarding the results of the S-H Resilience test, Group S had a lower score. Resilience is linked to academic success [19, 20]. Scores of STAI were higher in Group S. The higher State-Trait Anxiety score suggests students in Group S were in a stable anxiety state. The results of SF12v2 showed that students in Group S had lower scores in Role/Social component summary (RCS), in spite of their average scores in the Physical Functioning and Mental Health subscales. The low scores in Role/Social component summary (RCS) means that these students faced challenges pursuing a successful academic life.

Spectrum analysis of PRV indicated that those subjects who scored highly in LF/HF in Group S, which substitutes activities of the autonomic nerve, were inclined to hyper sympathetic status. HF, which was an indicator of parasympathetic activity, tended to be lower in Group S; even when we recorded the PRV in quiet conditions after five minutes of rest. Our findings may support the polyvagal theory by Porges, S.W. [21, 22] It is possible that some kind of intervention to adjust the autonomic nerve toward parasympathetic may improve the difficulties faced by the subjects in Group S. Non-linear analysis of accelerated plethysmography (LLE) showed little difference between Group S and Group H. LLE is a useful indicator of mental health [11]. A low level of LLE indicates that the subject is unable to adapt to external problems, which is characteristic of dementia and depression sufferers. A continuous high level of LLE implies external adaptability. Recently, the functioning brain network related to the state of rest was defined as the default mode network (DMN) [23, 24]. It is involved in the large scale brain network both during rest and cognitive tasks [25]. We suspect LLE may represent the brain activity and may have a relationship with DMN. We observed no difference in LLE and LLE (SD) between Group S and Group H. Our results for WAIS-III indicate that Group S had normal but lower scores in Working Memory Index (WMI) compared with Group H (Table 2). The presence of working memory deficits in high-functioning adolescents with ASD is disputed [26]. Group S had a lower score for working memory and had a lower performance score in the symbol search (Table 3). It suggests that visuospatial working memory functions were lower [27] compared with Group H.

Regarding the autoantibody against GAD65, we could not confirm the significance reported by Rout *et al.* [12] Recently, there have been reports of an association between ASD and anti-glutamate NMDA receptor antibodies [28, 29]. The etiology of ASD is thought to be multi-factorial. In further studies, we need to examine other autoantibodies against molecules related to brain function.

In conclusion, we found specific characteristics in Group S. They are as follows: 1) lower power of working memory despite high Full-Scale IQ, 2) higher ASD traits, 3) lower resilience power, 4) higher anxiety, 5) lower QOL in social health, 6) switching of autonomic nervous balance toward higher sympathetic activity. The results of our study revealed valuable information about the specificities of students who seek mental support. Interventions to improve these difficulties may contribute to improved academic achievement.

## Acknowledgments

We thank Tagami Y., Ikeda M., Tokunaga M., Miura T. and Ogasawara T. in Kanazawa University Health Service Center for their support to this study. We also thank Dr. Yamada M. and members of the Department of Neurology and Neurobiology of Ageing, Kanazawa University Graduate School of Medical Sciences for their support of our research activities.

This work was supported by JSPS KAKENHI Grant Number. 15H03084

